# GAP mimetic activity of pan-Ras TCI daraxonrasib synergizes with K-Ras Switch-II pocket inhibition

**DOI:** 10.64898/2026.03.11.711097

**Authors:** Patrick Pfaff, Kevan M. Shokat

## Abstract

Tricomplex inhibitors (TCIs) are a novel class of direct Ras inhibitors that target the GTP-bound Ras(on) state trough recruitment of Cyclophilin A. Daraxonrasib (RMC-6236) is a pan-Ras TCI that was recently shown to restore GTPase activity of G12-mutant Ras proteins. Structural analysis of a pan-Ras TCI bound to K-Ras(GDP– AlF_3_) reveals a transition-state arrangement of Tyr32 and Gln61 that closely resembles endogenous GTPase– GAP complexes. This includes a closed Switch-I conformation engaging the *cis*-GTPase machinery in a manner analogous to non-arginine-finger GAPs such as RanGAP. These observations position pan-Ras TCIs as pharmacologic GAP mimetics. The GTPase-promoting activity of daraxonrasib suggests synergy with Switch-II pocket K-Ras inhibitors, including the approved GDP-state selective K-Ras G12C inhibitor adagrasib (MRTX-849), whose engagement of K-Ras(GTP) is kinetically constrained by slow endogenous hydrolysis of the mutant GTPase. We demonstrate that daraxonrasib sensitizes K-Ras(GTP) to adagrasib labeling in both recombinant protein and cellular context. In K-Ras G12C and G12D mutant cell lines, combinations of daraxonrasib with adagrasib or HRS-4642 (MRTX-1133 analog) yield more rapid K-Ras engagement, rapid p-ERK suppression, and significant Loewe synergy scores in viability assays. These findings establish GAP mimetics as rational and potent combination partners for SW-II inhibitors. The synergistic combination has potential to deepen and prolong pathway suppression while enabling dose reductions that may mitigate on-target toxicity and resistance.

## Introduction

Small GTPases are a diverse family of protein switches that are tightly integrated in cellular signaling. In their GTP-bound ‘on-state’, GTPases are recognized by cellular effectors which elicit downstream signaling upon GTPase recruitment (McCormick and Wittinghofer, 1996; Wittinghofer and Nassar, 1996). Canonical GTPase activity results in hydrolysis of GTP and the resulting GDP-bound GTPases are generally considered to reside in an ‘off-state’. GTP-loading and GTP-hydrolysis are subject to regulation by cellular factors termed guanine exchange factors (GEFs) and GTPase-activating proteins (GAPs), respectively (Figure 1A, dashed lines) (Wittinghofer and Waldmann, 2000). This intricate signaling network is a common source of dysregulation in disease, most prominently emerging from oncogenic hotspot mutation of members of the Ras family. The most highly mutated isoform, K-Ras, has attracted particular interest as a drug target as mutation of Gly12 or Gln61 results in impairment of both intrinsic and GAP-activated GTP hydrolysis, thus enriching K-Ras in a constitutive GTP-bound ‘on-state’.

**Figure 1:**
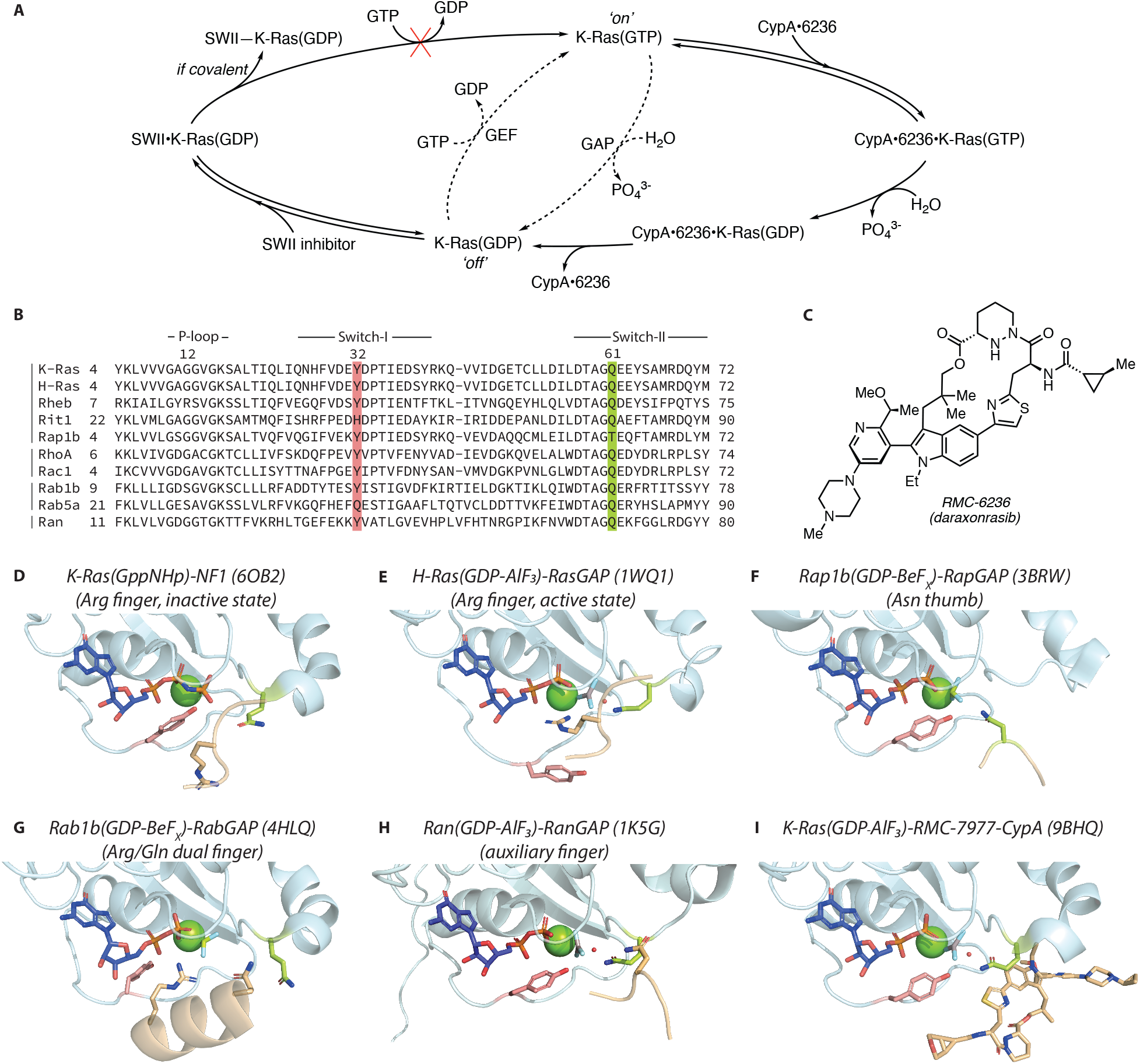
**A)** Regulation of K-Ras by endogenous GEF and GAP factors (dashed cycle) and pharmacologic engagement of K-Ras in either GDP- or GTP-bound state by SW-II inhibitors and daraxonrasib (RMC-6236), respectively (full cycle). **B)** Sequence alignment of small GTPases from diverse families (Ras, Rho, Rab and Ran), highlighting conserved Tyr32 and Gln61 in the Switch-I and -II regions. **C)** Chemical Structure of RMC-6236 (Daraxonrasib). **D)** K-Ras(GppNHp) bound to NF1 in an inactive Arg finger conformation (PDB: 6OB2). **E)** H-Ras(GDP-AlF_3_) bound to RasGAP in an active Arg finger conformation (PDB: 1WQ1). **F)** Rap1b(GDP-BeF_x_) bound to RapGAP in an active Asn thumb conformation; structure lacks catalytic water (PDB: 3BRW). **G)** Rab1b(GDP-BeF_x_) bound to RabGAP in an active Arg/Gln dual finger conformation; structure lacks catalytic water (PDB: 4HLQ). **H)** Ran(GDP-AlF_3_) bound to RanGAP with an auxiliary Asn finger but contributing its own catalytically active *cis*-Gln (PDB: 1K5G). **I)** K-Ras(GDP-AlF_3_) G12A bound to RMC-7977 (close structural analog of daraxonrasib) and CypA; CypA omitted for clarity (PDB: 9BHQ).

The discovery of covalent small molecules that can engage K-Ras G12C within a previously unknown cryptic pocket formed by the Switch-II region (SW-II pocket) (Ostrem et al., 2013) initiated a concerted effort between academia and industry towards targeting oncogenic K-Ras in the last decade (Ostrem et al., 2024). Initial efforts centered on the G12C-mutant state have led to regulatory approval of sotorasib (Canon et al., 2019; Lanman et al., 2020) and adagrasib (Fell et al., 2020; Hallin et al., 2020), with several next-generation candidates being investigated in the clinic. First-generation SW-II inhibitors display exclusive selectivity for the GDP-bound off-state (Janes et al., 2018; Patricelli et al., 2016). As a result, potent cellular trapping of K-Ras G12C relies on slow residual intrinsic GTPase activity and transient formation of the GDP state which is an isolated feature of the Cys mutant (Hunter et al., 2015). Mechanistically, engagement of the SW-II pocket locks K-Ras in its off-state due to blocking of GEF-induced nucleotide exchange (Figure 1A) (Ostrem et al., 2013). In addition, targeting of the G12C-mutant has informed the discovery and development of other mutant-specific SW-II pocket inhibitors. This includes the discovery of inhibitors for K-Ras G12D which has become tractable through both covalent (Zheng et al., 2024; Zheng and Shokat, 2025) and non-covalent engagement (Hallin et al., 2022; Vasta et al., 2022; Wang et al., 2022). In addition, chemical genetic experiments have demonstrated that the cryptic Switch-II pocket is a common feature of many GTPases beyond the Ras family which is targetable with similar chemical matter (Morstein et al., 2024).

In addition to SW-II inhibitors, another class of clinically promising drugs has recently emerged. In contrast to engaging Ras in the GDP-bound state, tricomplex inhibitors (TCIs) can specifically engage the GTP-bound ‘on-state’ of Ras proteins (K-Ras, H-Ras and N-Ras). TCIs recruit Cyclophilin A (CypA) to an interface formed by the Ras Switch-I and Switch-II regions which present high electrostatic complementarity with the CypA surface (Schulze et al., 2023). The resulting ternary complex sterically blocks effector binding resulting in high efficacy of TCIs. Similarly to the initial discovery of G12C-selective SW-II pocket inhibitors, TCIs were discovered through a G12C-targeted disulfide tethering screen of analogs of the natural product sanglifehrin A, a known natural product binder of CypA (Sanglier et al., 1999; Sedrani et al., 2003). These efforts have resulted in the discovery of mutant-specific covalent TCIs (G12C-specific elironrasib, RMC-6291 (Cregg et al., 2025b); G12D-specific zoldonrasib, RMC-9805 (Weller et al., 2025)) as well as a reversible pan-Ras TCI (daraxonrasib, RMC-6236 (Cregg et al., 2025a), Figure 1C) which are currently advancing in the clinic.

Strikingly, beyond simply blocking effectors from binding to GTP-bound Ras, it has been recently observed that the reversible pan-Ras inhibitor daraxonrasib and its close analog RMC-7977 can restore GTPase activity of mutant Ras (Cuevas-Navarro et al., 2025). This surprising finding implicates potential synergy between daraxonrasib and SW-II pocket inhibitors as enhanced turnover of K-Ras(GTP) to K-Ras(GDP) is expected to result in improved engagement of enriched K-Ras(GDP) by GDP-state selective SW-II inhibitors (Figure 1A). In this study, we explore the structural implications of daraxonrasib-induced GTPase turnover in the context of historical GAP structural biology across major GTPase families and present experimental evidence for synergy between pharmacologically restored mutant GTPase activity and GDP-state preferring Switch-II inhibition.

### Pharmacologic GAP mimetics in the context of endogenous GTPase-GAP pairs

The pioneering contributions of Alfred Wittinghofer’s laboratory have been transformative in elucidating how GTPase-activating proteins (GAP) accelerate GTP hydrolysis and in explaining how oncogenic mutations allow GTPases to escape GAP regulation (Wittinghofer et al., 1997). The determination of the crystal structure of Ras bound to the non-hydrolysable GTP analog GppNHp (Pai et al., 1990, 1989) revealed the conserved contacts of the γ-phosphate with the backbone atoms of Thr35 (Switch I) and Gly60 (Switch II), invariant residues across all GTPase families. The initial discovery of cytoplasmic RasGAPs (Adari et al., 1988; Trahey and McCormick, 1987) and an analogous role of the tumor suppressor NF1 (Ballester et al., 1990; Martin et al., 1990; Xu et al., 1990) by McCormick and others was followed by a period of speculation regarding the exact mechanism by which RasGAPs facilitate GTP hydrolysis. Wittinghofer’s group eventually demonstrated that RasGAPs utilize an arginine finger mechanism to accomplish an acceleration of up to 10_5_-fold, a catalytic rate at which dissociation of phosphate becomes rate limiting (Ahmadian et al., 1997b; Scheffzek et al., 1996). This was accomplished through systematic mutational experiments, combined with the use of aluminum fluoride as transition state analog (Mittal et al., 1996), mimicking the planar γ-phosphate transition state. Particularly insightful were kinetic and spectroscopic studies using neurofibromin (NF1), the physiological RasGAP mutated in neurofibromatosis type 1 patients, employing fluorescently labeled nucleotides, stopped-flow kinetics (Ahmadian et al., 1997a), and time-resolved Fourier-transform infrared (FTIR) spectroscopy using photocaged GTP (Cepus et al., 1998) performed collaboratively at the Max-Planck-Institute for Molecular Physiology.

These biochemical experiments were confirmed in detail by crystallographic studies which showed that RasGAP inserts an Arg finger into the Ras active site where it makes direct contacts with both the β- and γ-phosphates of GTP (Scheffzek et al., 1998, 1997), effectively neutralizing the resulting negative charges resulting from the GTPase reaction (Figure 1E). In addition, the backbone of the Arg finger contacts Gln61 of Ras which coordinates the catalytic water for attack of the γ-phosphate. The mechanism provided an explanation of how mutations at the Gly12 and Gln61 positions promote oncogenicity through increased Ras activity. Mutations of these essential residues result in impaired GTP hydrolysis given the direct involvement of Gln61 in catalysis and the placement of Gly12 in the middle of the active site, leading to steric clashes for any residue larger than Gly. Interestingly, in a structure that displays an inactive conformation of NF1 bound to Ras (Rabara et al., 2019) (Figure 1D), the catalytic Arg remains in a distal position to the active site and access to the γ-phosphate is blocked by Tyr32 (highlighted in red). Markedly, this setup is reversed in the active conformation of the RasGAP Arg finger (Figure 1E).

The Wittinghofer laboratory further had a central role in uncovering remarkable mechanistic GAP diversity among different GTPase families: while most Ras, Rho (Nassar et al., 1998; Rittinger et al., 1997) and Arf GTPases (Ismail et al., 2010) are subject to the archetypical Arg finger-dependent activation, other GTPases diverged from this paradigm. Rap1 lacks the highly conserved catalytic Gln61 present in other GTPases (Figure 1B, highlighted in yellow). Arg finger GAPs are thus unable to promote Rap1 GTP hydrolysis. It was discovered that RapGAPs instead provide an Asn thumb in *trans*, which replaces the function of the prototypical *cis*-Gln (Brinkmann et al., 2002; Daumke et al., 2004; Scrima et al., 2008) (Figure 1F). Similarly, Rheb, despite harboring an analogous Gln61 residue, has been shown to rely on a *trans*-Asn thumb provided by its cognate GAP TSC2 for catalytic turnover (Li et al., 2004; Yang et al., 2021), likely owing to steric blockade of the *cis*-machinery by a Rheb-characteristic Arg12 residue. Furthermore, Rab GTPases possess a conserved Gln61 residue yet also rely on a *trans*-acting residue to direct the catalytic water attack. In fact, most RabGAPs employ an Arg-Gln dual finger mechanism (Gavriljuk et al., 2012; Pan et al., 2006) (Figure 1G). Further, dual-specificity GAPs have been described with the fascinating ability to activate both Rap and Ras protein turnover by switching between Arg finger and Asn thumb mechanism depending on the substrate (Kupzig et al., 2006; Sot et al., 2010). Finally, RanGAP demonstrates a yet distinct mechanism by simply organizing the Switch-I and Switch-II regions of Ran in a catalytically competent conformation that relies on the prototypical Ran *cis*-Gln for coordinating the catalytic water attack (Bischoff et al., 1994; Seewald et al., 2002).

Interestingly, a common feature of such GTPase-GAP pairs not reliant on an Arg-finger, is a closed conformation of the switch I region, characterized by placement of Tyr32 within proximity of the γ-phosphate (cf. figures 1F and 1H). Tyr32 is a highly conserved residue among all families of small GTPases (Figure 1B, highlighted in red). Within the Ras family, Rit1 stands out for replacing Tyr32 with a His residue. Notably, no cognate GAP for Rit1 has been identified to date, and it has been speculated that Rit1 is mostly proteolytically regulated (Castel et al., 2019; Cuevas-Navarro et al., 2022). Similarly, some Rab family members such as Rab5a, do not possess Tyr32; it is likely a dispensable residue for Rabs due to the dual finger mechanism employed by RabGAPs and the requisite open conformation of Switch-I during GTP hydrolysis (Figure 1G).

The precise role of Tyr32 in promoting GTP hydrolysis has been debated but its involvement during intrinsic hydrolysis and in the highlighted GAP mechanisms proceeding from a closed Switch-I conformation appears likely (Cherfils et al., 1997). A few studies drawing from structural and biophysical experiments (Buhrman et al., 2010, 2007; Fink et al., 2024; Novelli et al., 2018) argue that Tyr32 may be involved in a water network generating a hydronium ion in the vicinity of the γ-phosphate, providing a positive charge similar to an inserted Arg finger, or assisting in proton transfer to the γ-phosphate within a bridging water network. However, these arguments are derived from ground state Ras structures bound to the non-hydrolysable nucleotide analog GppNHp, thus not allowing direct conclusions regarding the transition state of GTP hydrolysis.

In contrast to these models, a more recent study based on time-resolved crystallography following the intrinsic GTPase reaction of N-Ras by *in situ* cleavage of a photolabile GTP analog (Lin et al., 2025), a technique first utilized by Goody and co-workers (Schlichting et al., 1990, 1989), does not observe any excess water in the active site during the transition state of GTP hydrolysis. Tyr32 instead appears to be involved in coordination of the γ-phosphate and in displacement of excess water by promoting the closed Switch-I conformation, resulting in an entropically favorable transition state. A similar argument of favorable activation entropy through displacement of excess water has indeed been made in the case of Arg finger GAPs by Gerwert and Wittinghofer (Kötting et al., 2008), and thus Tyr32 coordination may fulfill a similar role in non-Arg finger GAP-mediated GTPase reactions. This is in line with simulation work favoring nucleophilic water attack directed by Gln61 from a sequestered active site during GTP hydrolysis, followed by rapid non-rate-limiting proton exchanges (Calixto et al., 2019). Interestingly, a Rap1a Y32F but not a Y32W mutant retained susceptibility to Asn-thumb RapGAP (Brinkmann et al., 2002), suggesting a dispensable role for the phenolic hydroxy group but highlighting the crucial role of a sterically compatible aromatic residue to produce a stable closed conformation of Switch-I. As an additional layer of Ras activity regulation, it has been observed that phosphorylation of Ras Tyr32 leads to RAF dissocation and RasGAP association, driven by electrostatic repulsion between the γ-phosphate and pTyr32 (Bunda et al., 2014; Kano et al., 2019); in line with the obligate open conformation of Switch-I required for Arg-finger GAP activity.

RMC-7977 is a close structural analog of daraxonrasib (RMC-6236) which has been reported to analogously induce GTP hydrolysis (Cuevas-Navarro et al., 2025). Strikingly, when examining the structure of RMC-7977 bound to K-Ras(GDP-AlF_3_) (Figure 5I, CypA excluded for clarity), Tyr32 and Gln61 adopt a highly analogous conformation to the transition state observed for Ran in complex with its auxiliary RanGAP (Seewald et al., 2002) (Figure 5H). This suggests a similar function of the pharmacologically induced ternary complex in orchestrating the GTPase reaction relative to the endogenous Ran GTPase-GAP pair. In addition, it was demonstrated that the GTPase promoting effect by RMC-7977 was abrogated for Gln61-mutant Ras, providing strong support for pharmacologic stimulation of the GTPase *cis*-machinery within the ternary complex. Ternary complex formation between Ras, CypA and RMC-7977 was strongly abolished for a K-Ras Y32S mutant in a control experiment. Yet, when examining the ternary structure, Tyr32 makes no direct contact within the ternary complex. This indicates an important role of Tyr32 in stabilizing the closed conformation required for competent ternary complex formation between Ras, CypA and TCI. The loss of binding upon Tyr32 mutation obscures inference on Tyr32’s role in catalytic turnover but given the close structural alignment in the transition state relative to the highlighted endogenous GTPase-GAP pairs encourages an important involvement in assisting the GTPase reaction. Indeed, when studying the original Ran-RanGAP pair, Wittinghofer and colleagues observed significantly reduced GAP-induction of GTPase activity for a Ran Y39A mutant, a site analogous to Ras Tyr32. In conclusion, this structural analysis reveals striking parallels between the catalytically competent daraxonrasib-CypA complex and endogenous GAP mechanisms highlighting daraxonrasib as a *GAP mimetic*. While Gln61 is a requisite for this current class of GAP mimetics due to utilization of the GTPase *cis*-machinery, alternative chemistry has been explored for pharmacologic induction of GTP hydrolysis in the absence of Gln61 (Zor et al., 1997; Ahmadian et al., 1999; Wang et al., 2025).

## Results

### Daraxonrasib displays synergy with Switch-II inhibition

Having examined the structural intricacies of restored GTPase activity within ternary complexes involving K-Ras(GTP), TCIs and CypA, we turned our attention towards studying the potential synergy between daraxonrasib and the clinically approved covalent G12C inhibitor adagrasib (MRTX-849, Figure 2A). As outlined in Figure 1A, we hypothesized that enhanced turnover of K-Ras(GTP) G12C to K-Ras(GDP) G12C would result in improved covalent engagement by GDP-state selective adagrasib. For an initial test of the hypothesis, we decided to study kinetic labeling of recombinant K-Ras G12C protein. Following EDTA-mediated nucleotide exchange, K-Ras G12C enriched in either GDP- or GTP-state was treated with adagrasib, and in the case of K-Ras(GTP) G12C the reaction was additionally conducted in the presence or absence of daraxonrasib or CypA, respectively. As expected, when incubating K-Ras(GDP) G12C with adagrasib, complete protein labeling was observed within minutes by liquid chromatography/mass spectrometry (LC/MS) (Figure 2B). K-Ras(GTP) G12C instead was engaged at a drastically reduced rate when treated with adagrasib alone. The retained partial engagement over time can be ascribed to slow residual intrinsic GTPase activity, a known effect that has been specifically observed for the K-Ras G12C mutant (Hunter et al., 2015). Similar observations were made when K-Ras(GTP) G12C was treated with either daraxonrasib (RMC-6236) or CypA alone, rendering each single agent incompetent of inducing potent GTP hydrolysis. However, when K-Ras(GTP) G12C was co-treated with daraxonrasib, CypA and adagrasib, significantly accelerated covalent engagement of K-Ras G12C was observed, in line with pharmacologically restored GTPase activity of the mutant protein, sensitizing K-Ras(GTP) G12C to covalent attack by GDP-state selective adagrasib.

**Figure 2:**
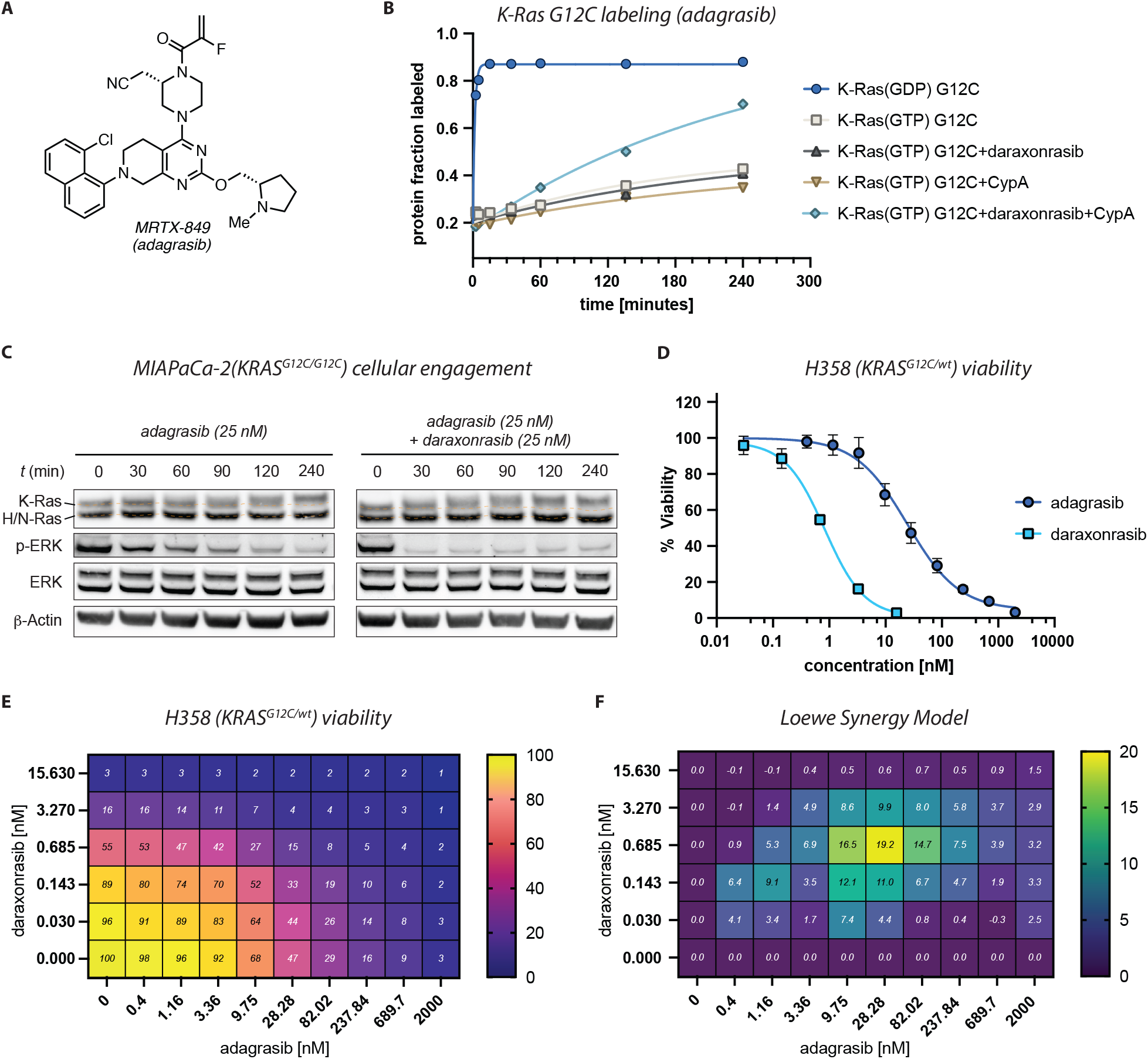
**A)** Chemical Structure of MRTX-849 (adagrasib). **B)** *In vitro* labeling experiment of recombinant K-Ras G12C CL protein (2 µM) by adagrasib (10 µM) in the presence or absence of daraxonrasib (RMC-6236, 5 µM) and CypA (5 µM), reactions were quenched at indicated timepoints and labeling was monitored by LC/MS. **C)** Western blot for cellular engagement of K-Ras G12C over time. MIAPaCa-2 cells were treated with adagrasib (25 nM) alone or in combination with daraxonrasib (25 nM) for the indicated periods. Cells were pre-treated with EGF (100 ng/mL) 15 minutes prior to drug addition. K-Ras G12C engagement was observed by mass band shift in Western blot, dashed parallel guidelines added to assist visualization of band shift in reference to unlabeled H/N-Ras. **D)** Cell viability of H358 cells (Cell Titer Glo) treated with adagrasib or daraxonrasib for 120 hours at the indicated concentrations. **E)** Cell viability of H358 cells (Cell Titer Glo) following combinatorial treatment at varying concentrations of adagrasib or daraxonrasib for 120 hours. **F)** Synergy scores calculated from viability observed in panel E using the Loewe Synergy Model (scores calculated using synergyfinderplus.org).

In a cellular context, mutant K-Ras G12C is maintained in a steady-state which is highly enriched in the GTP-bound state. We thus suspected that the enhanced labeling of K-Ras(GTP) G12C observed in vitro would also reflect in enhanced cellular engagement when combining adagrasib and daraxonrasib. To this end, MIAPaCa-2 cells, which harbor a homozygous *KRAS* G12C mutation, were treated either with adagrasib alone or with the combination of adagrasib and daraxonrasib for up to 4 hours and cell lysates were collected at the indicated timepoints (Figure 2C). To enrich K-Ras G12C uniformly in the GTP-loaded state, cells were pretreated with EGF 15 minutes prior to drug treatment. Lysates were then analyzed by Western blot to detect the extent of covalent K-Ras G12C engagement by mass band shift. Indeed, in cells treated with adagrasib alone, a clear K-Ras mass shift was not detectable until 120 minutes, with complete labeling achieved only by 240 minutes. In contrast, the combination treatment produced earlier target engagement, with labeling evident by around 60 minutes and full K-Ras labeling already observed at 90 minutes. This indicates accelerated daraxonrasib-induced turnover of cellular K-Ras(GTP) G12C to the GDP-bound state which can be potently engaged by adagrasib, in line with our observation for in vitro labeling of K-Ras(GTP) G12C (Figure 2B). In addition, when cells were treated with adagrasib alone, p-ERK signaling was reduced gradually over time in parallel with covalent K-Ras G12C engagement (Figure 2C), while the combinatorial treatment demonstrated very rapid p-ERK suppression, highlighting the dual role of daraxonrasib in both sensitizing the adagrasib attack and potently inhibiting downstream Ras signaling.

We next wondered whether the enhanced engagement of K-Ras G12C by adagrasib in the presence of daraxonrasib would also lead to a synergistic improvement of cellular potency with regard to cell viability. To this end, H358(*KRAS*_G12C/wt_) cells were treated at varying concentrations of adagrasib or daraxonrasib, either alone (Figure 2D) or in combination (Figure 2E). Examination of cell viability after a 120-hour treatment demonstrated enhanced potency of the combinatorial treatment at intermediate concentrations compared to either agent acting alone, a hallmark of synergy. Submission of this data to the Loewe synergy model (Figure 2F), resulted in a significant maximum synergy score of around 20 for a combination of roughly 40:1 of adagrasib to daraxonrasib, highlighting a significant role for daraxonrasib-induced sensitization of the G12C mutant cells towards adagrasib treatment.

Intriguingly, K-Ras G12D has been observed to possess the strongest sensitivity towards daraxonrasib-induced GTPase activity among K-Ras mutants (Cuevas-Navarro et al., 2025). While a GDP-AlF_3_ transition state-like structure of the G12D mutant in complex with a TCI and CypA has not been reported to date, it has been argued based on structures partially displaying transition state-like features that mutant Asp12 contributes favorably to coordinating the Gln61-dependent water network for nucleophilic attack of the γ-phosphate. While additional studies will be required to elucidate the enhanced GTPase activation for Ras G12D in greater detail, we suspected even stronger synergy for a combination of daraxonrasib with a K-Ras G12D-selective SW-II inhibitor in a G12D-mutant cellular context.

MRTX-1133 was the first selective, reversible K-Ras G12D inhibitor that was clinically explored (Hallin et al., 2022; Wang et al., 2022) (Figure 3A). Selectivity for the G12D mutant was achieved by engaging the K-Ras G12D Switch-II pocket through a salt bridge between mutant Asp12 and the bridged piperazine of MRTX-1133. Clinical investigation of MRTX-1133 has been halted with oral formulation issues cited by the parent company. However, a close structural analog of MRTX-1133, HRS-4642, featuring a fused oxazepane motif, continues to be clinically pursued as a liposome-formulated injectable therapy (Zhou et al., 2024) (Figure 3A). To this end, we decided to investigate the synergistic potential of daraxonrasib with HRS-4642 in AsPC-1 cells (*KRAS*_G12D/G12D_) (Figure 3B and C). Indeed, improved potency was observed when combining a low nanomolar concentration of HRS-4642 with a sub-nanomolar concentration of daraxonrasib. This resulted in an observed Loewe synergy score of above 30 for a ratio of 14:1 (HRS-4642 to daraxonrasib, Figure 3D). This improved synergy score suggests even stronger synergy for combining daraxonrasib in a G12D-mutant setting and is in line with the observation of enhanced hydrolytic turnover of K-Ras G12D compared to G12C.

**Figure 3:**
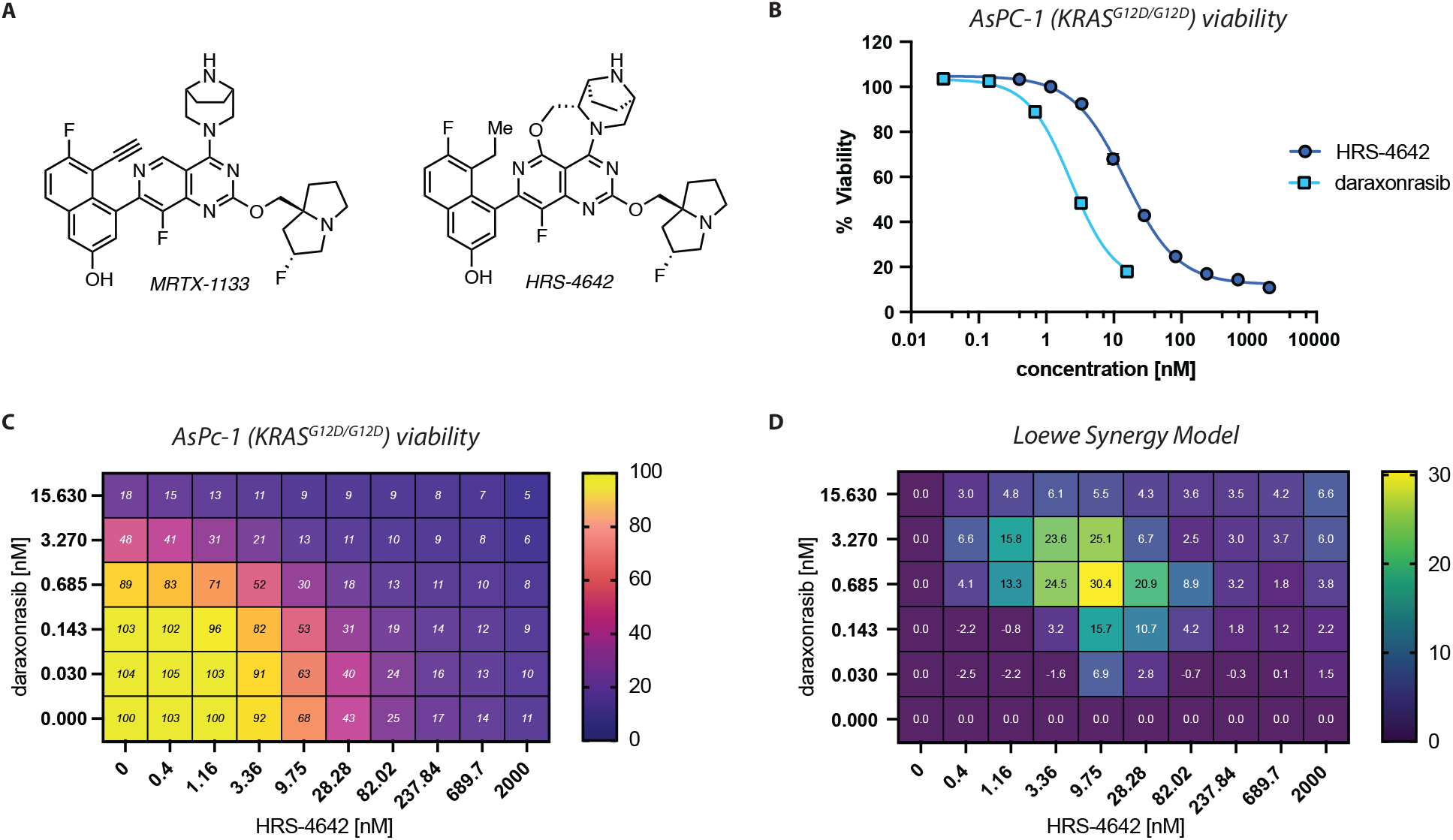
**A)** Chemical structures of MRTX-1133 and HRS-4642, respectively. **B)** Cell viability of AsPC-1 cells (Cell Titer Glo) treated with HRS-4642 or daraxonrasib for 120 hours at the indicated concentrations. **C)** Cell viability of AsPC-1 cells (Cell Titer Glo) following combinatorial treatment at varying concentrations of HRS-4642 or daraxonrasib for 120 hours. **D)** Synergy scores calculated from viability observed in panel C using the Loewe Synergy Model (scores calculated using synergyfinderplus.org).

## Discussion

K-Ras-mutant selective Switch-II pocket inhibitors and pan-Ras tricomplex inhibitors are two distinct modalities that have individually shown promise in addressing Ras-mutant cancer. Here we show that their combination demonstrates synergistic potential when compared to their use as single agents. Based on the recent observation that daraxonrasib restores GTPase activity of mutant Ras in complex with CypA (Cuevas-Navarro et al., 2025), we hypothesized that daraxonrasib would sensitize predominantly GTP-loaded mutant K-Ras to GDP-state selective Switch-II inhibition.

Our hypothesis for a catalytic role of daraxonrasib in enriching the adagrasib-sensitive GDP-loaded state was confirmed within in vitro labeling and cellular band shift assays for covalent engagement of K-Ras G12C. Our results suggest that daraxonrasib-induced hydrolytic turnover of Ras(GTP) G12C to Ras(GDP) G12C occurs at treatment-relevant timescales, providing additional turnover relative to residual innate GTPase or partially retained GAP activity (Li et al., 2021). Importantly, the high specificity of daraxonrasib-CypA for Ras(GTP) also implies rapid dissociation of the ternary complex upon hydrolysis, allowing engagement of the GDP state by Switch-II pocket agents further blocking GEF-driven reactivation of K-Ras. Indeed, the combination of daraxonrasib with either adagrasib or HRS-4642 demonstrated synergistic potency in K-Ras G12C and G12D mutant cell lines, respectively.

Additional roles of daraxonrasib beyond providing a kinetic advantage for Switch-II pocket occupancy are similarly likely. It has been shown that K-Ras-mutant cancer cells can partially adapt to specific inhibition of mutant alleles. A known adaptive mechanism is a lift of negative feedback on RTK expression including EGFR and HER2 resulting from rapid depletion of p-ERK upon acute mutant K-Ras inhibition (Feng et al., 2023; Ryan et al., 2020). The following receptor upregulation commonly initiates a rebound of p-ERK signaling through enhanced GTP-loading of wildtype Ras proteins including H-Ras and N-Ras. Indeed, combinations of SW-II G12C inhibitors with RTK blockade have been explored (Amodio et al., 2020; Yaeger et al., 2024) and are approved in colorectal cancer. We hypothesize that Switch-II pocket inhibition in combination with a low concentration of pan-Ras inhibitor can similarly intercept RTK-feedback-driven rewiring through additional inhibition of GTP-loaded wildtype Ras proteins (Holderfield et al., 2024); analogous effects have been observed when combining pan-Ras inhibitor RMC-7977 with G12C-specific on-state inhibitor RMC-4998 (Araujo et al., 2024).

In addition to adaptive responses, acquired resistance to mutant-specific Switch-II inhibition is common (Awad et al., 2021; Dilly et al., 2024; Tanaka et al., 2021). This includes secondary *cis*-alterations of K-Ras as well as amplification of the mutant allele, which increases the overall GTP-loaded protein fraction, a burden that can’t be overcome at maximum tolerable doses of Switch-II inhibitor. It has been shown that a mutant-specific Ras(on) inhibitor can assist in overcoming K-Ras(off) inhibitor resistance arising from enhanced K-Ras expression, albeit with low synergistic potential (Nokin et al., 2024). Given the high synergies observed here between pan-Ras(on) and mutant-specific K-Ras(off) inhibitors we anticipate that the pair holds even higher promise in resensitizing resistant tumor populations to mutant-specific Ras(off) inhibition. Similarly, orthogonal resistance mechanisms emerge for Ras(on) inhibition (Sang et al., 2025), and the highlighted combinatorial treatment may delay the onset of or increase the selection pressure prior to resistance.

Before the clinical investigation of daraxonrasib, hypothetic pan-Ras inhibition had been suspected to inflict systemic toxicity given the ubiquitous role of Ras in cellular signaling. Indeed, it was surprising when manageable safety profiles were reported within the ongoing clinical trials of daraxonrasib. This is likely facilitated through the GTP-state specific inhibition by tricomplex inhibitors allowing for cancer specificity (Wasko et al., 2024). However, on-target adverse events have been reported within ongoing clinical trials leading to dose modifications of daraxonrasib. Adagrasib can similarly produce treatment-related adverse events including hepatotoxicity (Chen et al., 2024). The synergy between daraxonrasib and adagrasib suggests that in combination either drug may be used at lower dosages than currently employed, thus potentially lowering systemic toxicity burdens.

GTPase Switch-II pocket inhibition remains a highly active field of research and next-generation agents including mutant-specific as well as pan-K-Ras modalities show promise as dual-state inhibitors for both GTP- and GDP-state. While the improved engagement of the GTP-bound K-Ras ‘on-state’ is an exciting feature, next-generation agents retain selectivity for the GDP-state (Maciag et al., 2025), likely owing to the higher flexibility and accessibility of the Switch-II pocket in the GDP-bound state. Accordingly, we expect the synergistic concept described here to persist for combinations of next-generation Switch-II inhibitors with Ras GTP hydrolyzers such as daraxonrasib.

Lastly, our structural analysis of the hydrolytically competent transition-state complex of K-Ras(GTP) bound by CypA and daraxonrasib revealed striking parallels with endogenous non-arginine finger GAPs, most notably the Ran-RanGAP system. This analogy is exemplified by the closed Switch-I conformation, with Tyr32 in contact to the γ-phosphate and Gln61 precisely aligned to direct the nucleophilic water attack, features central to the GTPase *cis*-machinery. These parallels underscore the pioneering groundwork of Alfred Wittinghofer’s laboratory in elucidating endogenous GTPase–GAP diversity and we anticipate these insights to be instrumental in the structure-guided development of next-generation pharmacologic GAP mimetics (Knox et al., 2024).

## Materials and Methods

### EDTA-mediated nucleotide exchange of recombinant protein

Recombinant K-Ras G12C cyslight protein (Morstein et al., 2024) was diluted to 4.0 µM in HEPES (20 mM, pH 7.5, 0.15 M NaCl, 1.0 mM MgCl_2_) and treated with EDTA (0.50 M stock, 10 mM final) and either GDP or GTP (50 mM stock, 0.40 mM final) for 2 hours at 4 ºC (total reaction volume 0.13 mL (GDP) and 0.50 mL (GTP), respectively). The nucleotide exchange reaction was quenched by addition of MgCl_2_ (2.0 M stock, 20 mM final). The quenched reaction mixture was desalted using a Zeba 7kDa MWCO spin column equilibrated in HEPES (20 mM, pH 7.5, 0.15 mM NaCl, 1.0 mM MgCl_2_) and the eluted nucleotide-exchanged protein was immediately used in the subsequent covalent labeling experiment.

### Intact protein mass spectrometry for covalent labeling of recombinant protein

12.5 µL of nucleotide-exchanged K-Ras G12C protein was distributed into 8 wells per time-course within a 384-well plate (4.0 µM stock, 2.0 µM final). All reactions were started simultaneously by addition of 12.5 µL of a 2X drug mixture (diluted in HEPES (20 mM, pH 7.5, 0.15 M NaCl, 1.0 mM MgCl_2_)) containing adagrasib (10 µM final) and/or daraxonrasib (5.0 µM final) and/or Cyclophilin A (5.0 µM final, Abcam ab86219), depending on the reaction (25 µL total reaction volume, 2% final DMSO). The individual reactions were quenched at the given timepoint by addition of 25 µL CH_3_CN/H_2_O containing 1.0% formic acid for a final protein concentration of 1.0 µM to be analyzed by LC/MS. The protein labeling reactions were analyzed by electrospray MS using a Waters Xevo G2-XS QTOF system following elution from an ACQUITY UPLC equipped with a ACQUITY Premier Protein BEH C4 300 Å 1.7 µm column. The mobile phase was a linear gradient of 5–95% acetonitrile/water containing 0.1% formic acid. The extent of protein modification was assessed following MaxEnt1 deconvolution (iterated to convergence at 1 Da resolution) of protein MS spectra.

### Cell culture

NCI-H358 (CRL-5807), MIAPaCa-2 (CRL-1420) and AsPC-1 (CRL-1682) cells were obtained from ATCC. NCI-H358 and MIAPaCa-2 cells were maintained in high-glucose DMEM (Gibco), supplemented with 10% heat-inactivated fetal bovine serum (FBS, Atlas Biologics). AsPC-1 cells were maintained in RPMI 1640 (Gibco), supplemented with 10% FBS. All cells and experiments were cultured at 37°C, 5% CO2.

### MIAPaCa-2 time course experiment

MIAPaCa-2 cells were seeded at 400k cells per well (6-well plate) to reach 40-60% confluency at the start of the experiment (12 to 24 hours post seeding). 15 minutes prior to drug treatment, the medium of the respective well was exchanged for 1.8 mL of fresh DMEM supplemented with 100 ng/mL recombinant EGF (PeproTech, AF-100-15). Cells were then treated at the indicated time prior to harvest with 0.20 mL of a 10X drug mixture (prepared in the same EGF-supplemented DMEM), containing either adagrasib alone (25 nM final) or adagrasib+daraxonrasib (25 nM final) (final DMSO concentration 0.2% for cell treatment). At the time of harvest, drug-containing medium was removed while cooling on ice. The cells were washed with ice-cold PBS (2.0 mL) and scraped in 40 µL RIPA buffer supplemented with protease and phosphatase inhibitors (mini cOmplete and phosSTOP, Roche). The lysates were transferred into tubes, flash frozen in liquid N_2_ and stored at -80 ºC until further processing.

### Gel electrophoresis and immunoblotting

Lysates were thawed on ice and clarified by high-speed centrifugation (>21000 rcf) for 10 minutes at 4 ºC. Total protein concentration of lysates was determined using protein BCA rapid gold assay and adjusted to 2.0 mg/mL. Normalized samples were mixed with 4x NuPAGE LDS sample buffer (Invitrogen) supplemented with 10x NuPAGE sample reducing agent (500 mM DTT) and heated at 70 ºC for 10 minutes.

SDS-PAGE was performed using NuPAGE 12% Bis-Tris gel (Invitrogen) in XT MES running buffer (Bio-Rad) at 200V for 90 minutes while cooling on ice. Proteins were transferred onto 0.2 μM nitrocellulose membranes using a semidry transfer device (Invitrogen iBlot 3). Membranes were blocked using Intercept TBS Blocking Buffer (LICOR) for 15 minutes at room temperature. Primary antibody incubation was performed in the presence of the indicated antibodies diluted in Intercept antibody diluent (LICOR) for at least 24 hours at 4 ºC. The membranes were washed four times with TBST (1 short wash, 3 x 5-minute wash) and blots were incubated with secondary antibodies (goat anti-rabbit IgG-IRDye800, goat anti-mouse IgG-IRDye 680, 1:5000 dilution in Intercept antibody diluent, LICOR) for 1 hour at room temperature. The membranes were washed four times with TBST (1 short wash, 3 x 5-minute wash) and imaged on a ChemiDoc MP imaging system (Bio-Rad).

### Primary antibodies and dilutions

> pan-Ras (Abcam, 108602, 1:1000)
>
> β-Actin (Cell Signaling Technology, 3700, 1:1000)
>
> ERK (Cell Signaling Technology, 4695, 1:1000)
>
> p-ERK(T202/Y204) (Cell Signaling Technology, 9101, 1:1000)

### Cell viability assay

Cells (H358 or AsPC-1) were seeded at 2k cells per well in the 60 inner wells of a 96-well plate (Greiner Cellstar µCLEAR®, WHITE, PS, F-Bottom, 655098) in 100 µL of appropriate medium (DMEM or RPMI, 10% FBS, respectively), and the 36 outer wells were filled with 200 µL PBS. Following incubation overnight (12-24 hours), cells were treated in technical triplicate with the indicated compounds at the indicated concentration using a D300e digital dispenser (Tecan) at a normalized concentration of 0.1% DMSO. Cells were incubated for 120 hours at 37 ºC, 5% CO_2_. Cells were allowed to cool to room temperature for 30 minutes prior to addition of 20 µL Cell-Titer Glo reagent (Promega) per well. Plates were incubated at room temperature on an orbital shaker for 15 minutes prior to detection of luminescence on a Clariostar Plus platereader (BMG Labtech).

## Notes

K.M.S. has consulting agreements for the following companies, which involve monetary and/or stock compensation: BridGene Biosciences, Erasca, Exai, G Protein Therapeutics, Genentech, Kumquat Biosciences, Kura Oncology, Lyterian, Merck, Montara Therapeutics, Nextech, Revolution Medicines, Pfizer, Rezo, Tahoe, Totus, Type6 Therapeutics, Wellspring Biosciences (Araxes Pharma)

## Acknowledgments

P.P. was supported by a postdoctoral fellowship from the Swiss National Science Foundation (SNF Mobility Grant P500PN_214278). K.M.S. thanks the Sjöberg Foundation for supporting this work.

